# Monocyte Subset Predicts Clinical Outcomes in Fibrotic Hypersensitivity Pneumonitis

**DOI:** 10.1101/2025.09.25.678673

**Authors:** Sonia M. Leach, Brian Vestal, Evans R. Fernández Pérez

## Abstract

**Rationale:** Hypersensitivity pneumonitis (HP) is an immunologically mediated form of lung disease resulting from inhalational exposure to various antigens. While there is growing understanding of the immune cells involved in HP, the relationship between clinical outcomes and circulating cell types in HP is not well established.

**Methods:** Single-cell RNA sequencing was performed on peripheral blood mononuclear cells (PBMCs) from 40 patients with fibrotic hypersensitivity pneumonitis (fHP) in a clinical trial of pirfenidone (NCT02958917). After quality control, we compared data from 33 fHP patients with 36 sex-matched healthy controls (GSE196735). Using Seurat v5, we harmonized the data, formed clusters, and reduced dimensionality through UMAP. Cell type identities were assigned based on a published reference (GSE271789), with clusters annotated for predominant cell type(s) present in at least 30% of the cells. We compared cell type proportions between fHP and HC using the chi-squared test and performed differential expression analysis with DESeq2 on pseudobulk counts, adjusting for sex. Genes significantly upregulated in fHP versus HC at an FDR-adjusted p-value of 0.01 were used to create a multivariate predictive signature for clinical outcomes in fHP participants. Performance was assessed through leave-one-out cross-validation for prediction accuracy.

**Results:** Compared to HC, fHP has an increased proportion of non-classical CD16+ monocytes (4.2% fHP vs. 1.5% HC), C14+ monocytes with a myeloid-derived dendritic phenotype (2.1% fHP vs. 0.1% HC), platelets (1.0% fHP vs. 0.3% HC), plasmacytoid dendritic cells (0.5% fHP vs. 0.1% HC) and predominantly classical CD14+ monocytes (29.5% fHP vs. 3.4% HC, adjusted-p<0.001). In patients with fHP, cell clusters were associated with the presence of baseline chest CT honeycombing, increased extent of CT lung fibrosis, and worse progression-free survival. CD14+ monocytes consistently demonstrated a high prediction accuracy (>0.90) for these clinical metrics.

**Conclusions:** Peripheral monocyte clusters may be valuable prognostic markers, potentially helping identify at-risk fHP patients.

## Background

Hypersensitivity pneumonitis (HP) is an immunologically mediated complex lung disease resulting from inhalational exposure to a large variety of inciting antigens. A subset of patients developed progressive pulmonary fibrosis (fibrotic HP, fHP), a leading cause of excess morbidity and mortality.

Research using bulk RNA sequencing of peripheral blood mononuclear cells (PBMCs) from fHP patients at initial presentation identified a 74-gene signature distinguishing progressive from non-progressive fHP,^1^ suggesting that the PBMC transcriptome may be informative of immune mechanisms in fHP. However, the specific immune cell types involved remained unclear. Recently, single-cell transcriptomics in fHP patients uncovered novel immune perturbations.^2^ Yet, the relationship between circulating cell types and clinical outcomes has not been assessed. Furthermore, there are no studies evaluating the association between immune cell abnormalities and disease severity using quantitative CT that provide a reproducible assessment of lung fibrosis extent. Therefore, we aimed to determine the prognostic value of cell-specific clusters by conducting single-cell RNA sequencing (scRNAseq) on PBMC samples from a prospective cohort of fHP patients.

## Methods

### Study population

Samples were obtained from 40 adult fHP patients enrolled in the 52-week pirfenidone randomized clinical trial (NCT02958917).^3^ Demographics, pulmonary function and CT lung fibrosis extent quantified by data-driven texture analysis (DTA)^3,4^ were collected at the time of blood draw. Clinical outcomes include the annualized changed in FVC and progression-free survival (PFS) defined as the time from randomization to the first occurrence of either: 1) A relative decline of ≥10% in FVC and/or DLCO, 2) Acute respiratory exacerbation, 3) A decrease of ≥50 meters in the 6-minute walk distance, 4) Increase in background prednisone by ≥10 mg or introduction of corticosteroids and/or steroid-sparing therapy because of clinical deterioration, or 5) Death.

After quality control, data from 33 fHP patients were compared to 36 sex-matched healthy controls (Gene Expression Omnibus Accession GSE196735, pools 1-3). Standard analysis using Seurat v5 in R v4.1.1 removed cells with fewer than 200 expressed genes and greater than 15% mitochondrial reads, normalized within each sample by scTransform, integrated samples with Harmony, and used 30 principal components to calculate the Uniform Manifold Approximation and Projection (UMAP) and Louvain clusters at resolution 0.8.

### Statistical analysis

Cell type identities were assigned by projection onto a published reference (GSE271789), and each cluster was annotated to the cell type(s) occurring among 30% or more of the cells in the cluster. Comparing fHP and healthy controls per cluster, differences in cell type proportions were determined by the chi-squared test for homogeneity with Bonferroni tests post-hoc, and differential expression was analyzed using DESeq2 on pseudobulk counts, adjusting for sex. Genes significantly upregulated in fHP versus healthy controls at a false discovery rate (FDR)-adjusted p-value=0.01 in each cluster were then used to construct multi-variate predictive signatures for four clinical outcomes among the fHP participants.

Using custom R scripts, the outcome was first binarized, if necessary, using a percentile threshold. DESeq2 tests were used to remove genes associated with sex and then to determine candidate genes with univariate associations with the outcome. Elastic net (EN) regularization was performed using the glmnet R package on the set of candidate univariate gene features, using 50 trials of balanced, random subsets of the data, where class size was balanced to that of the minority class. If the number of features selected in >50% of the EN trials exceeded five, we then used all samples to perform multivariate partial least-squares discriminant analysis (PLS-DA) with one latent factor using the mdatools R package.

We applied step-forward variable selection among the set of candidate features using leave-one-out cross-validation of prediction accuracy. We report the performance and feature set for the PLS-DA model with maximum cross-validated prediction accuracy. We considered outcomes for baseline honeycombing (binary), progression-free-survival (PFS, binary), annualized FVC change (25th percentile), and baseline DTA (30th percentile). Pathway analysis was conducted using EnrichR in R.

## Results

The HC participants’ age ranged from 19 to 97, with 73% being older than 60 years. At baseline, fHP patients had a mean age of 66±5.9 years, a mean FVC% of 59±11% and a mean DLCO% of 52.8±16.1. Their 6-minute walk distance was 393.0±98.0 meters. The DTA score was 32.6±15, with 13 (39%) showing CT honeycombing. Twenty-two (66%) were on pirfenidone. Fifteen patients had BAL, and 93% (14/15) had a macrophage-to-lymphocyte ratio >1.

Compared to HC (**Figure 1**), fHP has an increased proporpotion of non-classical CD16+ monocytes (4.2% fHP vs. 1.5% HC, adjusted-p<0.001), classical CD14+ monocytes (29.5% fHP vs. 3.4% HC, adjusted-p<0.001), C14+ monocytes with a myeloid-derived dendritic phenotype (2.1% fHP vs. 0.1% HC, adjusted-p<00.1), platelets (1.0% fHP vs. 0.3% HC, adjusted-p<00.1) and plasmacytoid dendritic cells (0.5% fHP vs. 0.1% HC, adjusted-p<00.1). Among patients with fHP, these cell clusters were associated with baseline disease severity and progression (**Figure 1**). Notably, among all cell clusters, CD14+ monocytes consistently demonstrated a high prediction accuracy (>0.90) across all these clinical metrics.

**Figure 1.**
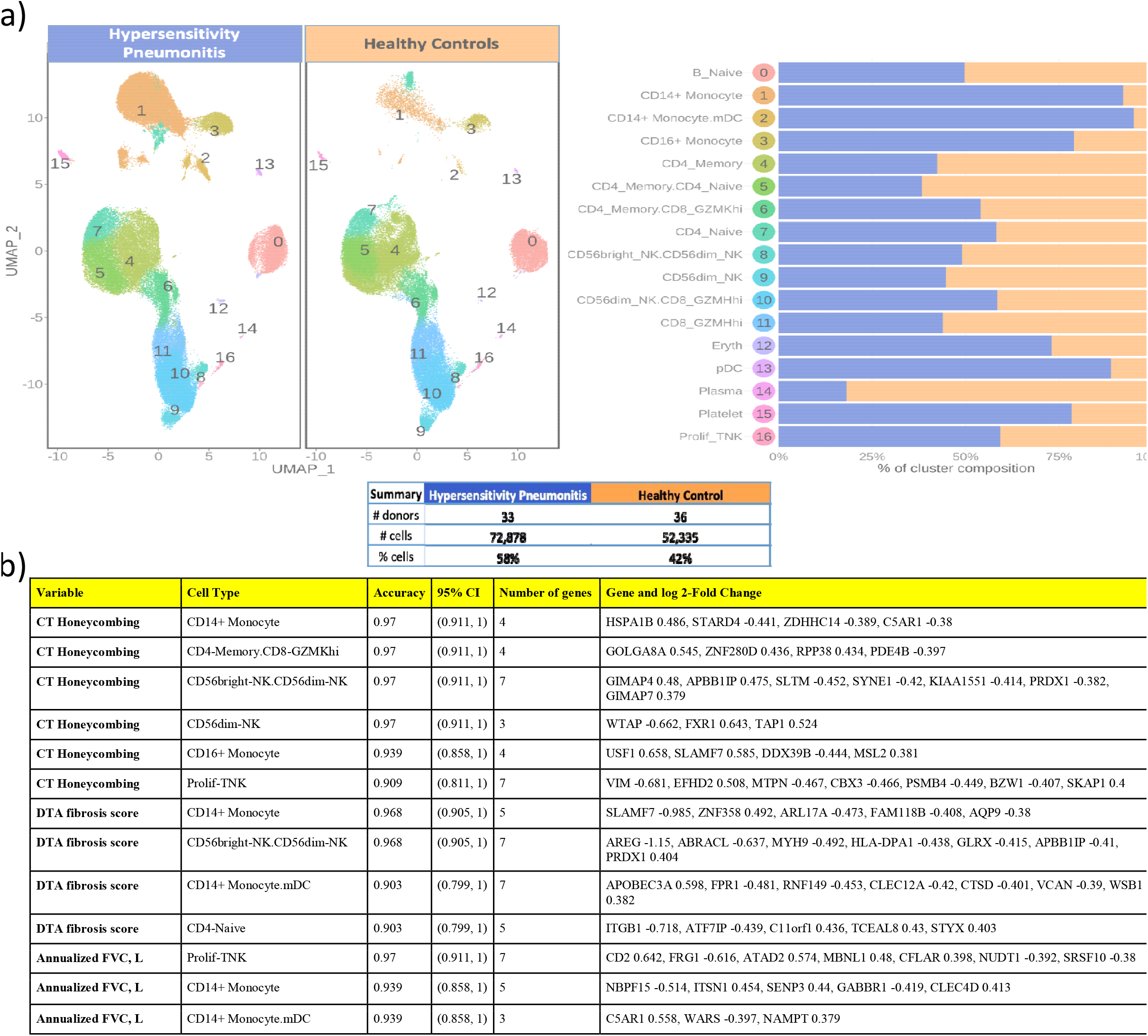
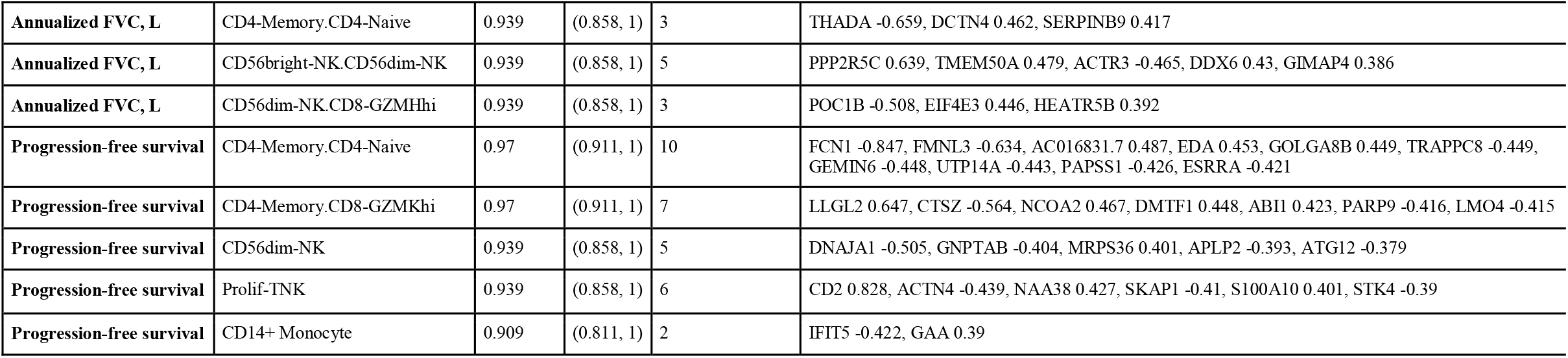
Single cell data in fibrotic HP: **a)** Immune cell populations in PBMCs in fibrotic HP (blue) vs. healthy controls (orange). **b)** Predictive signature accuracy in fibrotic HP for clinical measures. **Abbreviations: DTA:** data-driven texture analysis. **FVC:** forced vital capacity. **DLCO:** diffusing capacity of the lungs for carbon monoxide. **PFS:** progression free survival defined as the time from study treatment randomization to the first occurrence of any of the following events: A relative decline of ≥10% in FVC% or DLCO%; Acute respiratory exacerbation; A decrease of 50 meters or more in the 6 minute walk distance; Death. **30p:** below 30th percentile. **25p:** below 25th percentile.

### Pathway analysis revealed that CD14+ monocytes associated with

- The presence of CT honeycombing is highly enriched in the complement cascade (C5AR1), intracellular cholesterol homeostasis (STARD4), and protein palmitoylation (ZDHHC14), all implicated in CD14+ monocyte fibroinflammatory responses.^5,6^
- High DTA score is highly enriched with cellular permeability and metabolism (AQP9), vesicular trafficking and lysosomal function (ARL17A), transcriptional regulation in response to cellular stress (FAM118B), and cell-to-cell communication (SLAMF7), all of which also play an intricate role in monocyte-driven inflammation.^7,8^
- Lower annual FVC is highly enriched with pathogen recognition (CLEC4D),^9^ cell activation (ITSN1),^10^ and inflammatory signaling and stress responses (SENP3).^11^ They are critical regulators of innate immune cell functions and their dysregulation may contribute to the development of a pro-inflammatory and pro-fibrotic environment.
- Increased PFS is highly enriched by interferon signaling and modulation of NF-κB signaling (IFIT5), implicated in aberrant wound-healing response,^12^ autoimmune diseases^13^ and cancer.^14^

## Discussion

Our findings suggest that peripheral blood monocytes, particularly intermediate monocytes (CD14+CD16+), monocyte-derived dendritic cells, and predominantly classical monocytes (CD14+CD16-), are associated with disease severity and clinical outcome in fHP patients. In a lung microenvironment marked by injury and inadequate resolution, these monocytes, recruited from the bloodstream to lung tissue, can serve as precursors of polarized pro-fibrotic macrophages,^15^ contributing fHP progression.

This study extends previous work on immune abnormalities in fHP.^2^ Compared to controls, fHP patients showed reduced peripheral memory B-cells and CD56^hi^ NK-cells. However, we also found reduced CD4+ memory and CD8 GZMH^hi^ cells, likely due to trafficking to the lungs.

Our results underscore the need to better understand monocytes and other cell cluster patterns, along with their transcriptional activity, to enhance novel molecular-clinical approaches for classifying HP that may more accurately predict clinically meaningful outcomes.

Our study has limitations, including the absence of a validation cohort. Currently, no existing fHP cohort integrates peripheral blood single-cell measurements with either disease severity through computer-based methods (radiomics) and longitudinal follow-up. Nonetheless, our findings align with previous work involving 39 fHP patients,^2^ suggesting potential generalizability. To mitigate biases due to small sample size and class size imbalance limitations, we prefiltered sex-associated genes, used balanced class bootstrapping during modelling, and applied leave-one-out cross-validation. Also, bulk RNA-seq analysis demonstrated no significant effect of pirfenidone, so all participants were used in this analysis. Lastly, although we were unable to confirm specific cell clusters in lung tissue, most fHP patients with BAL displayed an elevated macrophage-to-lymphocyte ratio, consistent with studies indicating an enrichment of macrophages in lung fibrosis.

In summary, our analysis of peripheral blood scRNAseq reveals significant differences in the immune landscape of fHP compared to HC. This study suggests that monocyte clusters may be valuable prognostic markers, potentially helping identify at-risk fHP patients and guiding treatment decisions.

